# First report of Infectious Spleen and Kidney Necrosis Virus in Nile tilapia in Brazil

**DOI:** 10.1101/2020.10.08.331991

**Authors:** Henrique César Pereira Figueiredo, Guilherme Campos Tavares, Fernanda Alves Dorella, Júlio César Câmara Rosa, Sóstenes Apolo Correia Marcelino, Felipe Pierezan

## Abstract

In June 2020, an atypical fatal outbreak in the Nile tilapia farm was investigated. Twenty-three animals were collected and different tissues were used for bacterial isolation, histopathological and electron microscopic examination, and viral detection using molecular methods. A large number of megalocytes were observed in the histopathological analysis of several tissues. Icosahedral virions, with a diameter of approximately 160 nm, were visualized inside the megalocytes through transmission electron microscopy of the spleen tissue. The virions were confirmed to be *infectious spleen and kidney necrosis virus* (ISKNV) through PCR and sequencing analyses of the fish samples. This viral pathogen is fatal in the early stages of Nile tilapia farms in the USA, Thailand, and Ghana; however, until now, there have been no reports from Brazil. In 14 of the sampled fish, *Streptococcus agalactiae, Edwardsiella tarda*, or *Aeromonas hydrophila infections* were also detected. This is the first report of fatal ISKNV infections in the Brazilian Nile tilapia fish farms.

## 1. Introduction

Nile tilapia (*Oreochromis niloticus*) is a key fish species for freshwater aquaculture worldwide, with a global production estimated at 4,525,400 tons (FAO, 2020). In Brazil, Nile tilapia is the majorly cultured fish species (approximately 758,000 tons), placing the country as the fourth largest producer in the world (PeixeBR, 2020).

This fish species is susceptible to several bacterial pathogens such as *Streptococcus agalactiae* (Mian et al., 2009), *Francisella orientalis* (Leal et al., 2014), *Aeromonas hydrophila* (Nicholson et al., 2020), and *Edwarsiella tarda* (Niu et al., 2019); as well as some viral agents such as tilapia lake virus (Eyngor et al., 2014), betanodavirus (Bigarré et al., 2009), and iridovirus (Subramaniam et al., 2016). However, no viral infections have been reported in Brazil to date.

The infectious spleen and kidney necrosis virus (ISKNV) is a member of the genus *Megalocytivirus* belonging to the family Iridoviridae. This virus has been associated with systemic disease and high mortality in many freshwater and marine fish species in aquaculture (Wang et al., 2007; Subramaniam et al., 2012), mostly in Asian countries. The first report of this pathogen infecting Nile tilapia was from Thailand in 2015 (Dong et al., 2015) and USA in 2016 (Subramanian et. al., 2016). In both reports, outbreaks were observed in tilapia fry and juveniles, with clinical manifestations of lethargy, gill pallor, and ascites, and a mortality rate of about 75%. Recently, a mass mortality associated with ISKNV infection was described in Ghana, in cage culture systems of Nile tilapia in Lake Volta (Ramírez-Paredez et al., 2020).

In this study, we report the first case of ISKNV infection in Nile tilapia in Brazil, based on the molecular analysis, histopathology, and electron microscopic observations. Furthermore, some epidemiological aspects of the outbreak and co-infections with bacterial pathogens have been described.

## 2. Materials and Methods

In June 2020, a fatal outbreak in the Nile tilapia farm located in Gouvelândia municipality (Geographic coordinates: 18°38’31″ S, 50°03 °’57″ W), Goias State, Brazil, was investigated. The fish were acquired from commercial larviculture (approximately 25 g). They had already been vaccinated against *Streptococcus agalactiae* (Aquavac Strep Sa ™) and were reared in floating cages installed in the São Simão reservoir (722.25 km^2^ and volume of 5.54 billion m^3^). After a period of approximately two months of growth, the fish were subjected to classification management. A few days later, mortalities were observed in three lots, with the maximum mortality rate reaching 6.4%. The water temperature in this period was 24.5 ± 1.5°C. During the growth period, there was no introduction of new batches of fingerlings in the farm. A total of 23 diseased fish (average weight of 261.52 g ± 185.93) were sampled, transported on ice, and received in the laboratory for analysis.

Swabs of the brain, kidney, and spleen were aseptically obtained from these fish, streaked onto 5% horse blood agar (HBA) or cysteine heart agar (CHA) supplemented with 1% bovine hemoglobin, and incubated at 28°C for 2–7 days to isolate bacterial pathogens. Bacterial species were identified using matrix-assisted laser desorption-ionization time-of-flight (MALDI-TOF) mass spectrometry with a Bruker Microflex MALDI Biotyper 2.0 (Bruker Daltonics, USA), as previously described (Assis et al., 2017).

Fragments of the brain, kidney, spleen, liver, heart, gill, intestine, and stomach from four fishes were collected for histological analysis. Of these tissues, only those from the liver and spleen were subjected to transmission electron microscopic analysis. In addition, the brain, kidney, and spleen tissues were also collected and pooled for the molecular diagnosis of ISKNV, tilapia lake virus (TiLV), and betanodavirus (VNNV – viral nervous necrosis virus).

For histological analysis, tissues were fixed in neutral buffered formalin (NBF) for 48 h prior to processing using standard protocols in a vacuum infiltration processor (Leica ASP300S). Tissue sections were embedded in paraffin and stained with hematoxylin and eosin (H&E) using a standard histological procedure. Stained sections were examined for general histopathology using light microscopy (Olympus BX41) and images were recorded using an automated upright microscope system (Leica DM4000 B).

For electron microscopy, small samples of the spleen and liver, fixed in NBF as above, were rinsed three times with 0.1 M sodium cacodylate buffer, followed by post fixation in 2% glutaraldehyde in the same buffer prior to a second post fixation for 1 h in 1% osmium tetroxide in 0.1 M sodium cacodylate buffer. Subsequently, the fixed tissues were contrasted with uranyl acetate (2% in deionized water), dehydrated in an ascending series of ethanol solutions, embedded in Epon resin, and sectioned (60 nm). The sections were collected on uncoated copper grids and contrasted with lead citrate. The sections were viewed using a Tecnai-G2-12-Spirit electron microscope at 120 kV (FEI, USA), and digital images were recorded.

For molecular diagnosis, the collected tissues were subjected to total DNA and RNA extraction using the Maxwell® 16 Tissue DNA Purification Kit and Maxwell®16 LEV simplyRNA Tissue Kit, respectively, in the Maxwell 16 Research Instrument (Promega, USA) following the manufacturer’s instructions. Extracted nucleic acids were quantified using a Nanodrop spectrophotometer (Thermo Scientific, USA) and stored at −80°C until use. Extracted DNA or RNA were used as the template for detection of Red Sea Bream Iridoviral Disease (RSIVD) / ISKNV using PCR, and VNNV and TiLV using quantitative reverse-transcription PCR (RT-qPCR).

The following primers 1-F: 5′-CTCAAACACTCTGGCTCATC-3′ and 1-R: 5′-GCACCAACACATCTCCTATC-3′ were used for the amplification of the gene sequence (570 bp) of both RSIV and ISKNV. PCR assays were performed as previously described (Kurita et al., 1998) with some modifications. Reactions were performed with a HotStart Taq polymerase kit (Qiagen, USA) in a final reaction volume of 25 μL. The reaction mixture consisted of 1× PCR buffer, 1.0 μM of each PCR primer, 0.2 μM dNTPs, 1.25 U Taq DNA polymerase, and 50 ng template DNA. The PCR conditions were as follows: an initial denaturation at 95°C for 15 min, followed by 30 cycles of 94°C for 30 s, 58°C for 1 min, and 72°C for 1 min; final elongation was carried out at 72°C for 5 min. The primers used were purchased from Integrated DNA Technologies (IDT, Coralville, USA). pRSIVD plasmid (GeneArt, Germany), containing a 570 bp target regions of the primers, was used as positive control. The Veriti 96-Well Thermal Cycler was used and amplicons were separated in the QIAxcell Advanced using QX DNA Screening Kit (Qiagen).

Detection of VNNV and TiLV was performed using the Promega GoTaq Probe 1-step RT-qPCR System and both were performed in the QuantStudio 7 (Applied Biosystems, UK) as follows: 30 min at 45°C, 2 min at 95°C and 40 cycles of 15 sec at 95°C and 1 min at 60°C. The VNNV reaction was performed in a final reaction volume of 20 μL. The mix composed of 1× GoTaq® Probe qPCR Master Mix (2X), 0.9 μM of each PCR primer (RNA2 FOR: 5′-CAACTGACARCGAHCACAC-3′; RNA2 REV: 5′-CCCACCAYTTGGCVAC-3′); 0.75 μM of probe (RNA2 probe: 5′-FAM-TYCARGCRACTCGTGGTGCVG-BHQ1-3′) (Panzarin et al., 2010), 1x GoScript™ RT Mix for 1-Step RT-qPCR (50X), and 100 ng template RNA. TiLV detection was performed using the primers TiLV-93F 5′-AGCCTGCCACACAGAAG-3′; TiLV-93R 5′-CTGCTTGAGTTGTGCTTCT-3′ and probe TiLV-93Probe 5′-FAM-CTCTACCAGCTAGTGCCCCA-IowaBlack-3′ (Waiyamitra et al., 2018). The reaction mix consisted of 1× GoTaq® Probe qPCR Master Mix (2X), 0.6 μM of each PCR primer (TiLV-93F and TiLV-93R), 0.3 μM of probe (TiLV-93Probe), 1x GoScript™ RT Mix for 1-Step RT-qPCR (50X), and 100 ng template RNA. pVNNV and pTiLV plasmids were used as positive controls.

In case of positive PCR results for ISKNV, sequencing of the PCR products was carried out. The PCR amplicons were purified using AMPure XP da Agencourt (Beckman Coulter) according to the manufacturer’s instructions. Sequencing reactions were performed using a BigDye™ Terminator Cycle Sequencing Kit (Applied Biosystems) and evaluated with an ABI 3500 Genetic Analyzer (Life Technologies, USA). Forward and reverse sequencing products were used to generate contigs using the BioEdit software (Ibis Biosciences, USA) version 7.2. Identity was evaluated using the BLAST webserver (http://www.ncbi.nlm.nih.gov/BLAST) by checking against existing sequences in the nt/nr database.

## 3. Results

The clinical signs observed in the sampled fish were anorexia, melanosis, mucus hypersecretion, integument hemorrhage, ascites (Figure 1a), and gill pallor (Figure 1b). At necropsy, hepatomegaly (Figure 1c), splenomegaly, hemorrhagic viscera (Figure 1c), friable muscle, presence of liquid in the celomatic cavity (Figure 1d) were observed, and in some fish, the viscera were liquefied, especially the liver and spleen. Bacterial isolates were obtained from the brain, kidney, or spleen from 14 out of 23 diseased Nile tilapia (Table 1). Among these isolates, *Streptococcus agalactiae* (*n* = 6 fish), *Edwardsiella tarda* (*n* = 7), *and Aeromonas hydrophila* (*n* = 2) were identified using MALDI-TOF analysis. In total, 9 out of 23 fish presented negative results for the bacteria. In molecular diagnosis, all fish were positive for RSVID/ISKNV in PCR analysis. PCR amplification yielded a product with an average size of 539 bp. The sequences obtained had a query coverage and identity greater than 99% and 99.24%, respectively, when compared to other sequences deposited in public database (NCBI) by BLAST analysis. For all sequences, the greatest similarity observed was with infectious spleen and kidney necrosis virus isolate NBFGR:ISKNV:C5 laminin-like protein gene (Accession number: MT224131.1), confirming that the fish were infected with ISKNV (Table 1). There was no detection of VNNV and TiLV in the samples. Co-infection with bacteria such as ISKNV + *S. agalactiae* in six of the sampled fishes; ISKNV + *E. tarda*, in six; ISKNV + *A. hydrophila*, in one; and ISKNV + *E. tarda* + *A. hydrophila*, in one, were detected (Table 1).

**Table 1.**
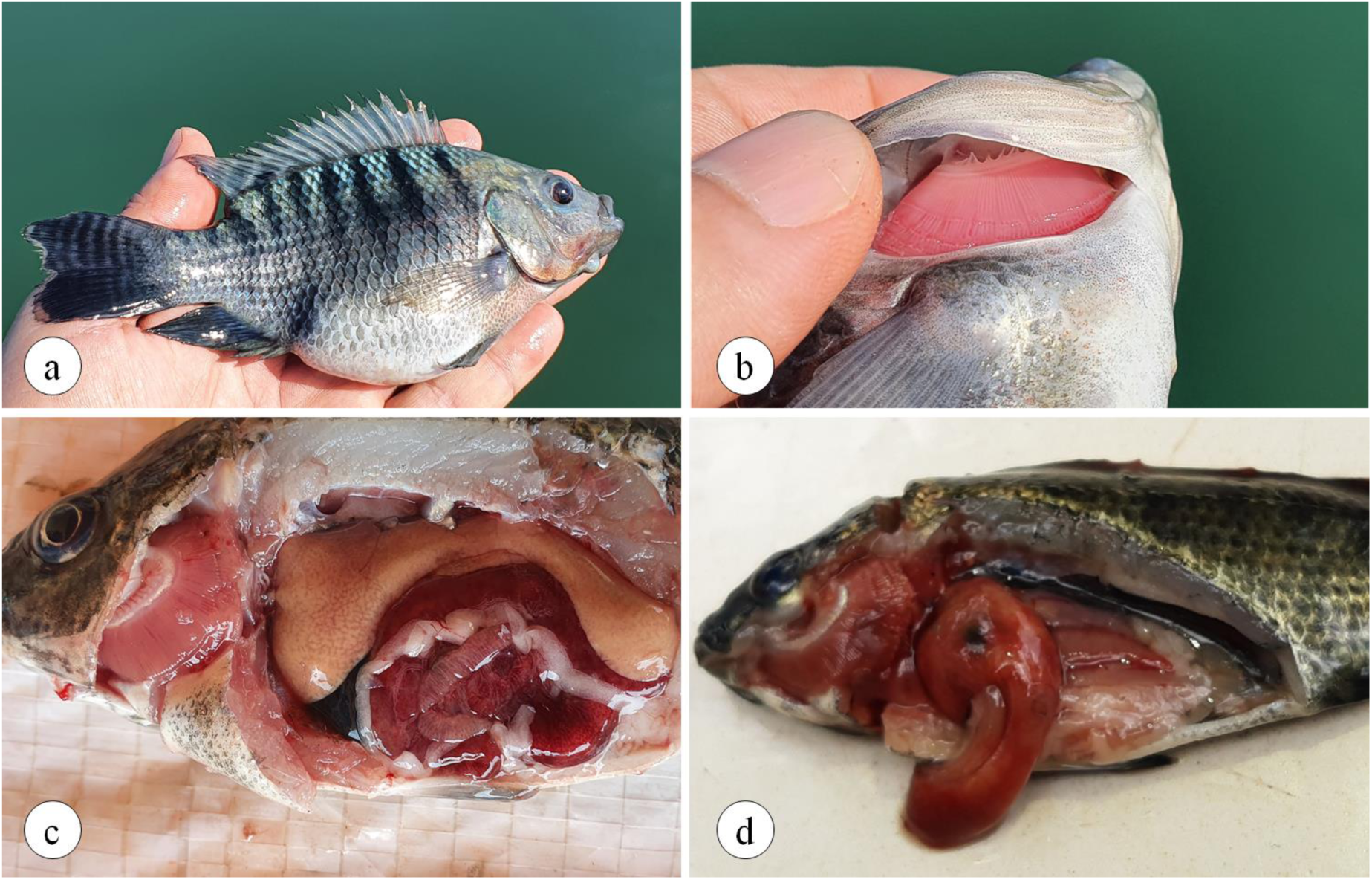
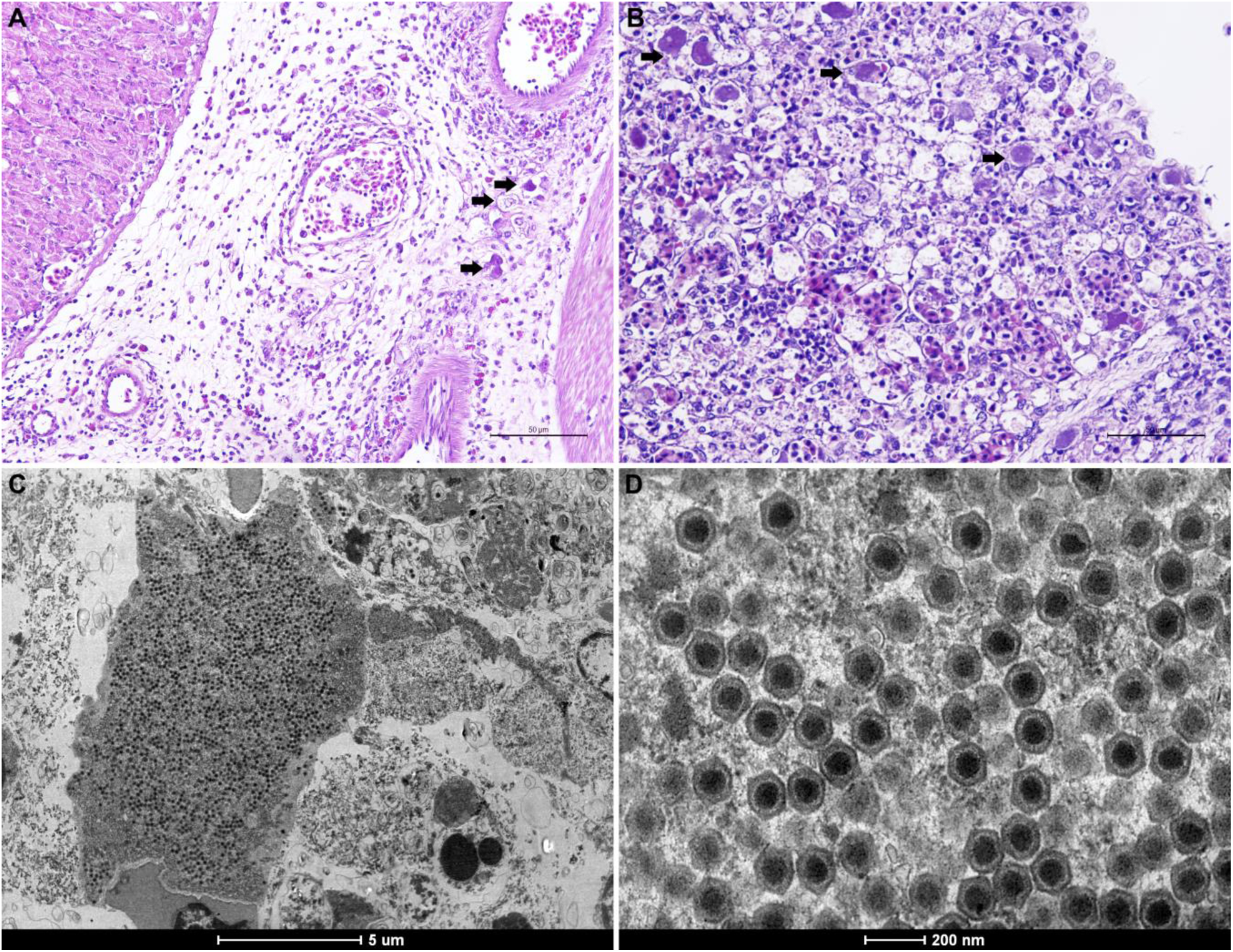
Information about the animals collected, ID, weight, tissue sampled, bacteriology, and molecular diagnosis results.

**Figure 1.**
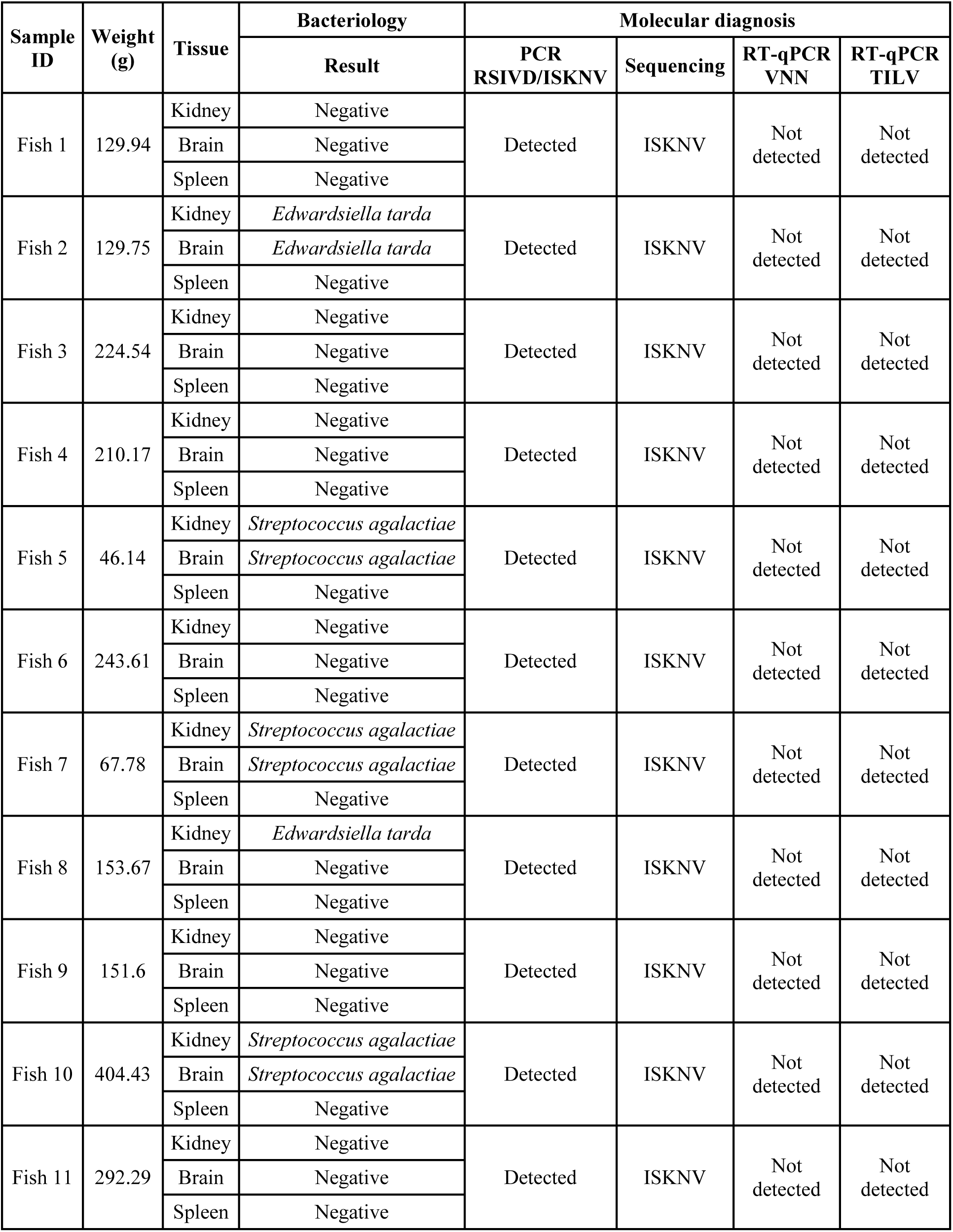
Diseased Nile tilapia with ascites (a), gill pallor (b), hepatomegaly and hemorrhagic viscera (c) and the presence of liquid on the celomatic cavity and liver congestion (d)

In histopathological analysis, several tissues presented basophilic (Figure 2AB), hypertrophied cells (megalocytes), often in large numbers in the anterior kidney and splenic parenchyma, and in smaller numbers in the hepatic and cardiac parenchyma, glomerulus of the posterior kidney, gills, and the intestinal and gastric submucosa. These cells were often situated near blood vessels and were larger in size (20–30 μm in diameter) with a pale foamy or intensely basophilic granular appearance. Degenerative features such as necrosis, and hemorrhagic foci were often seen in the splenic, hepatic, and renal parenchyma. Loss of the renal glomeruli and cells in the intestinal and gastric submucosa were observed; however, the renal tubules were usually unaffected. Accumulation of mononuclear cells was observed in a few instances in the heart and around blood vessels in the liver.

**Figure 2.**
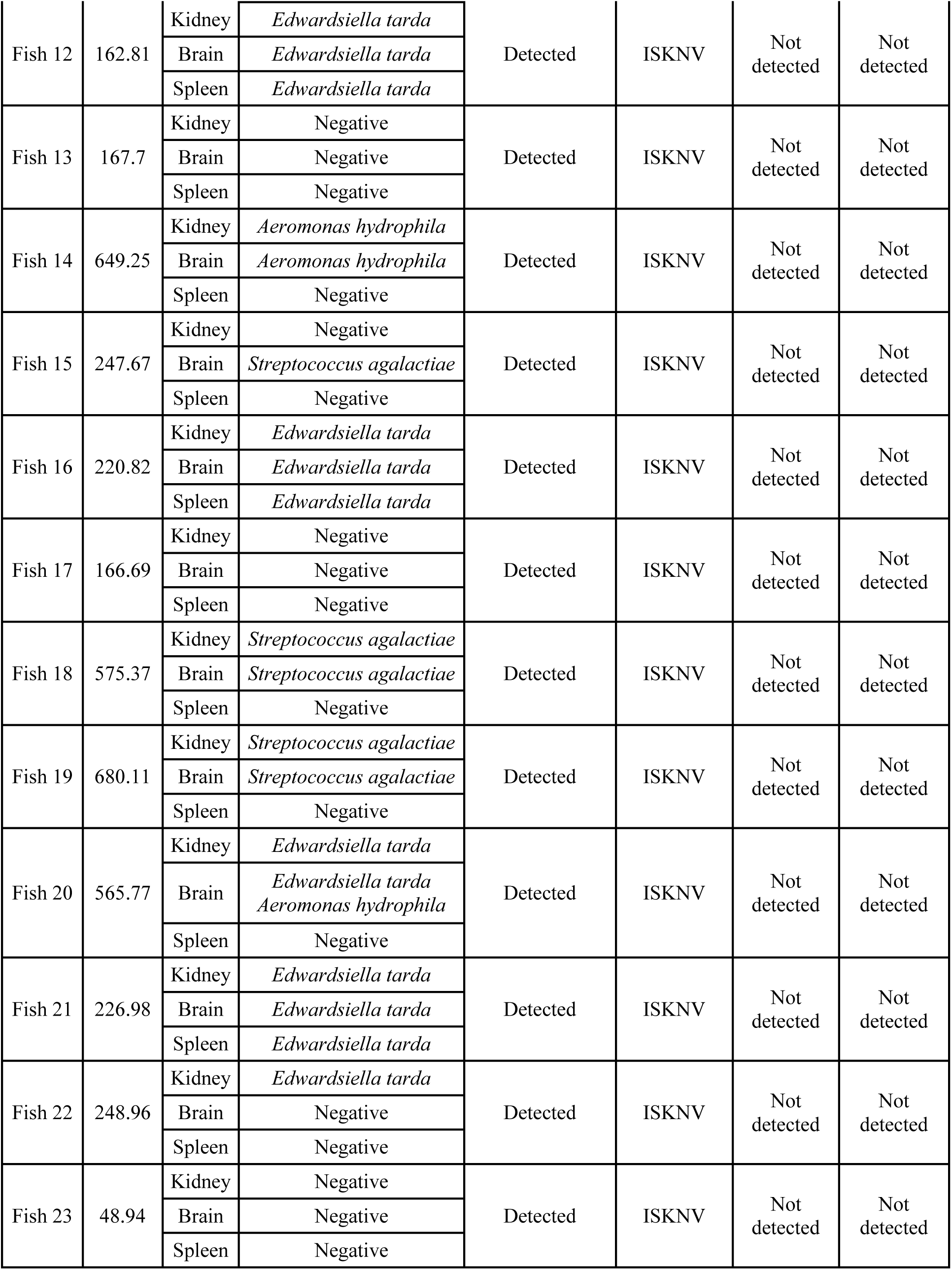
Histopathology and electron microscopy of organs of Nile tilapia infected with ISKNV. A. Section of the stomach of an infected fry showing basophilic hypertrophied cells (arrow) located at the periphery of blood vessels of the submucosa. Bar = 50 μm. B. Section of the spleen of an infected fry showing basophilic hypertrophied cells (arrow) scattered throughout the parenchyma. Bar=50 μm. C. Electron micrograph of an individual infected cell with numerous viral particles. Scale bar = 5 μm. D. Numerous lightly stained icosahedral virions showing details of the outer capsid and inner membrane and central electron dense core. Scale bar = 200 μm

Numerous cells containing electron-dense bodies in the cytoplasm were observed in the spleen (Figure 2C) during transmission electron microscopy analyses. These cells had numerous diffusely spread virions, which formed ‘arrays’. Virions were icosahedral and approximately 160 nm in diameter with an external double membrane and a central electron dense core (Figure 2D).

## 4. Discussion

In this study, ISKNV was detected in diseased Nile tilapia from a Brazilian farm. In previous ISKNV outbreaks in fish, the viral pathogen was associated with massive mortalities in the fry and juvenile stages (Subramaniam et al., 2016; Ramírez-Paredez et al., 2020). This was also observed in our investigation. The clinical signs and pathological alterations observed after necropsy are also consistent with these reports. In addition to Nile tilapia, ISKNV has been associated with mortality in farmed barramundi (Dong et al., 2017), bluegill sunfish (Liu et al., 2019), mandarin fish (He et al., 2000), Murray cod (Go et al., 2006), and ornamental fish species (Tanaka et al., 2014; Jung-Schroers et al., 2016). Therefore, ISKNV is considered as an emerging pathogen in many countries.

In Brazil, ISKNV infection has only been reported in ornamental fish (Maganha et al., 2018); however, atypical mortalities in juvenile Nile tilapia have been reported by fish farmers since 2019, without a definitive diagnosis. We received diseased fish (juveniles with an average weight of 261.52 g) from one of these fish farmers, and performed an investigation using bacteriological, histopathological, and molecular diagnosis assays. All sampled animals were positive for ISKNV as observed through PCR and sequencing techniques. The histopathological and electron microscopic findings were similar to those described in other studies (Subramaniam et al., 2016; Ramírez-Paredez et al., 2020). The presence of megalocytic cells in many tissues and the visualization of viral particles in the spleen confirm ISKNV as one of the etiological agents of this fatal outbreak. However, bacteriological examination demonstrated co-infection of the virus and bacteria in the same fish. *E. tarda, S. agalactiae, and A. hydrophila* were the key bacteria isolated from diseased fish. Natural co-infection of bacterial pathogens and ISKNV in diseased Nile tilapia has already been described in other studies (Dong et al., 2015; Ramírez-Paredez et al., 2020) and it seems to be a common feature of this viral infection. The impact of these co-infections on the severity of the outbreak as well as the possible control measures to be adopted need to be addressed.

Although this viral pathogen belongs to the same genus as the causative agent of Red Sea Bream Iridoviral Disease (RSIVD), *Megalocytivirus* (Kurita and Nakajima, 2012), the disease is not listed by World Organisation for Animal Health (OIE) as a notifiable disease from aquatic animals. Nevertheless, as it is an exotic pathogen in Brazilian tilapia farms, we have communicated the existence of a new viral disease in the country to the National Official Veterinary Service.

In this study, we could not confirm the origin of the infection because clinical signs and mortality were observed only two months after introduction of the fish into the production system. Although other fish farmers shared the same aquatic environment as the affected farm, being part of the same reservoir, there were no reports of fish transport among farms, and fry acquisition from another source. This may suggest the indirect transmission of the viral pathogen via water. On the other hand, the clinical manifestation of the disease after fish classification may indicate that the animals were carriers of the virus, and that handling acted as a pre-disposing factor.

Robust studies are necessary to determine the occurrence of ISKNV in other tilapia farms and its impact on the national production of this aquatic species. There is the possibility of the infection being transmitted horizontally (He et al., 2002) and vertically (Suebsing et al., 2016), as well as the possibility of the pathogen being present in the carrier fish (Subramaniam et al., 2012). Therefore, the establishing a surveillance program and monitoring health in hatcheries and broodstock is recommended to avoid the transmission of the disease among farms.

In conclusion, to our knowledge, this is the first report of a fatal ISKNV outbreak in Nile tilapia farms in Brazil as well as the description of the co-infection of this viral pathogen with some bacteria such as *S. agalactiae* and *E. tarda*. Taken together, our findings represent a new challenge to the commercial production of Nile tilapia in the country.

## Acknowledgements

The authors would like to acknowledge the Center of Microscopy at the Federal University of Minas Gerais for providing the equipment and technical support for electron microscopy analyses. This study was supported by grants of FAPEMIG and CNPq.

